# Dissemination and Stakeholder Engagement Practices Among Dissemination & Implementation Scientists: Results from an Online Survey

**DOI:** 10.1101/627042

**Authors:** Christopher E. Knoepke, M. Pilar Ingle, Daniel D. Matlock, Ross C. Brownson, Russell E. Glasgow

## Abstract

**Introduction:** There has been an increasing focus on disseminating research findings, but less about practices specific to disseminating and engaging non-researchers. The present project sought to describe dissemination practices and engagement of stakeholders among dissemination & implementation (D&I) scientists.

**Methods:** Methods to disseminate to and engage non-research stakeholders were assessed using an online survey sent to a broad, diverse sample of D&I scientists.

**Results:** Surveys were received from 210 participants. The majority of respondents were from university or research settings in the U.S. (69%) or Canada (13%), representing a mix of clinical (28%) and community settings (34%). 26% had received formal training in D&I. Respondents indicated routinely engaging in a variety of dissemination-related activities, with academic journal publications (88%), conference presentations (86%), and reports to funders (74%) being the most frequent. Journal publication was identified as the most impactful on respondents’ careers (94%), but face-to-face meetings with stakeholders were rated as most impactful on practice or policy (40%). Stakeholder involvement in research was common, with clinical and community-based researchers engaging stakeholder groups in broadly similar ways, but with critical differences noted between researchers with greater seniority, those with more D&I training, those based in the US, and those in community vs, clinical research settings.

**Conclusions:** There have been increases in stakeholder engagement, but few other practices since the 2012 survey, and some differences across subgroups. Methods to engage different stakeholders deserve more in-depth investigation. D&I researchers report substantial misalignment of incentives and behaviors related to dissemination to non-research audiences.

## Background

Dissemination, defined as “an active approach of spreading evidence-based interventions to the target audience via predetermined channels using planned strategies” (2), is the critical process linking research findings to practitioners who can implement them, leading to benefits among the people or communities of interest. Frequently recommended dissemination practices to reach non-research audiences include “Designing for Dissemination” (3, 4), use of multiple channels, development of guides to program implementation, and engagement of multiple types of stakeholders in the development and evaluation of interventions and dissemination plans. The number of publications on dissemination has increased dramatically over the years (5–7) since classic work on diffusion of innovations (8). What is less known is the extent to which there have been increases in the use of evidence based and best practices 1 among dissemination and implementation (D&I) researchers, and if there are differences in 1 dissemination practices across different types of D&I researchers.

Differences in preferred sources of information between researchers and practitioners have been documented and researchers are increasingly urged to “go beyond” academic publication and presentations at major professional conferences (3, 9). It is not known more specifically what avenues and strategies researchers, and especially D&I researchers, use to facilitate translation of their findings into practice and policy.

There has been a strong encouragement to meaningfully engage patients and community stakeholders in research from PCORI, NIH, and other organizations (10). Two relatively recent developments of interest have been use of social media and stakeholder engagement practices (11, 12). While each of these has existed for decades, most health care and public health researchers have not been early adopters of these approaches, we were interested in what specific engagement strategies D&I researchers use and the extent to which they used them.

The use of a variety of dissemination practices to non-research audiences (e.g., publication, meetings, webinars) has previously been described by Brownson et al. (2013). They surveyed a sample of public health researchers in 2012 concerning their dissemination practices, including which dissemination practices the researchers rated as most impactful, and which were most aligned with incentives for career advancement. This survey provided valuable information, but was seven years ago and we hypothesize that there have been significant increases since then due to trends encouraging dissemination, due in part to the number of newly trained D&I scientists (13–18). Additionally, the 2012 survey did not extensively assess practices such as designing for dissemination or stakeholder engagement, nor did it include clinical researchers or non-U.S. researchers. At that time, few researchers had received formal D&I training. Thus, an updated and expanded assessment of current dissemination practices was warranted.

The purposes of this current project were to: 1) conduct a survey conceptually similar to the Brownson et al. 2012 survey by characterizing current practices of research dissemination to non-research audiences among D&I researchers; 2) include a broader sample of D&I scientists; 3) include additional dissemination and stakeholder engagement practices; and 4) investigate potential researcher characteristics associated with greater use of various dissemination strategies.

## Methods

### Survey development

The survey was developed by beginning with the Brownson et al. and Tabak et al. surveys (1, 19). We adopted and, in many cases, modified questions and/or response options in this survey to address 2018 priorities, evolution of the field, and a greater number of dissemination practices. We also added several items related to investigator characteristics and stakeholder engagement practices. The primary domains assessed included dissemination practices; impressions of the impact and importance of different practices for a) impact and b) promotion; stakeholder engagement practices; and respondent characteristics. Due to the different and expanded sample, we also needed to modify several items (e.g., to address medical as well as public health researches; Canadian and other researchers in addition to U.S. researchers). Initial drafts of the survey were iteratively developed and refined by team members, reviewed by original 2012 researchers, and reactions from members of key target audiences. We also deleted several items to keep the survey to a reasonable length. A copy of the survey, which is publicly available for others to use, is presented in Appendix A.

### Sampling frame

Participants were recruited to take part in an online survey assessing self-reported practices related to dissemination of findings to non-research audiences, as well as methods by which respondents engage stakeholders in research to enhance translation. Potential survey respondents were identified by having productivity or recent training related to D&I science. Participants were recruited through a variety of organizations, lists, and avenues (see Table 1). As described later, it was not possible to determine a denominator of scientists invited or an accurate return rate because: a) several organizations did not allow us access to mailing lists or to send individual e-mails so we did not know the number of invitations sent, b) the number of incorrect e-mails, or c) the degree of overlap among the different sampling sources. We expect that the latter was very large due to the number of scientists who may well have been funded by D&I funders, published in *Implementation Science*, and been trained in D&I research. The survey was powered through Qualtrics online survey software.

**Table 1:** Sampling Frame by Source

| Table 1: Sampling Frame by Source |  |  |
| --- | --- | --- |
| Sample Description | Time Frame Included | Approximate N invited |
| Corresponding authors in *Implementation Science* | January 2015-December 2017 | 225 |
| PCORI grantees with D&I foci | 2015- 2017 | 111 |
| CIHR KT grantees | 2015-2017 | 55 |
| IRI fellows | 2011-2017 | 63 |
| MT-DIRC fellows | 2014-2017 | 55 |
| CDC Prevention Resource Center leaders | 2017 | 47 |
| VA QUERI listserv | April 2018 | Unknown |
| KT Canada listserv | May 2018 | Unknown |
| NCI D&I listserv | April 2018 | Unknown |
| TIDRH listserv | April 2018 | 313 |

\*Sample n's do not account for likely overlap between sources

### Survey implementation

Surveys were distributed through Qualtrics (when individual e-mail addresses were available) or through electronic listservs as appropriate. Listserv distributions were conducted by managers of those listservs rather than by our study personnel due to confidentiality requirements. Potential participants for whom we had individual e-mail addresses received up to three reminder emails at one-week intervals from April-May 2018. Responses were collected anonymously and respondents did not receive any incentive for participation. The project was approved by the local Institutional Review Board, including a waiver of written consent.

### Analyses

Primary analyses followed an *a priori* analytic plan, consisting primarily of descriptive statistics, percentages, frequencies, and narrative comparisons. In instances when the analytic plan called for subgroup comparisons, independent samples T-test, chi squares, and chi square likelihood ratio tests were used as appropriate. Finally, logistic regression analyses were performed to evaluate independent contributions of several potential respondent characteristic predictors of use of high levels of stakeholder engagement. A priori predictions were that: 1) the 2018 sample would report greater use of dissemination practices in addition to the usual publications and presentation strategies than the 2012 sample; 2) those receiving formal D&I research training would engage in more dissemination practices and more stakeholder engagement activities than those not; and 2) that Canadian researchers would make greater use of stakeholder engagement practices than U.S. researchers. The majority of remaining analyses were descriptive and exploratory in nature.

## Results

### Respondents

Surveys were received from 210 total participants, 68% (148) employed by U.S.-based universities, conducting research in a number of contexts and with a variety of training and professional experiences (Table 2). While we were able to determine that no single individual completed the survey more than once, we were unable to calculate a response rate due to 1) the unknown denominator information for listservs, and 2) an unknown quantity of individuals who were likely in two or more of our target groups.

**Table 2:**
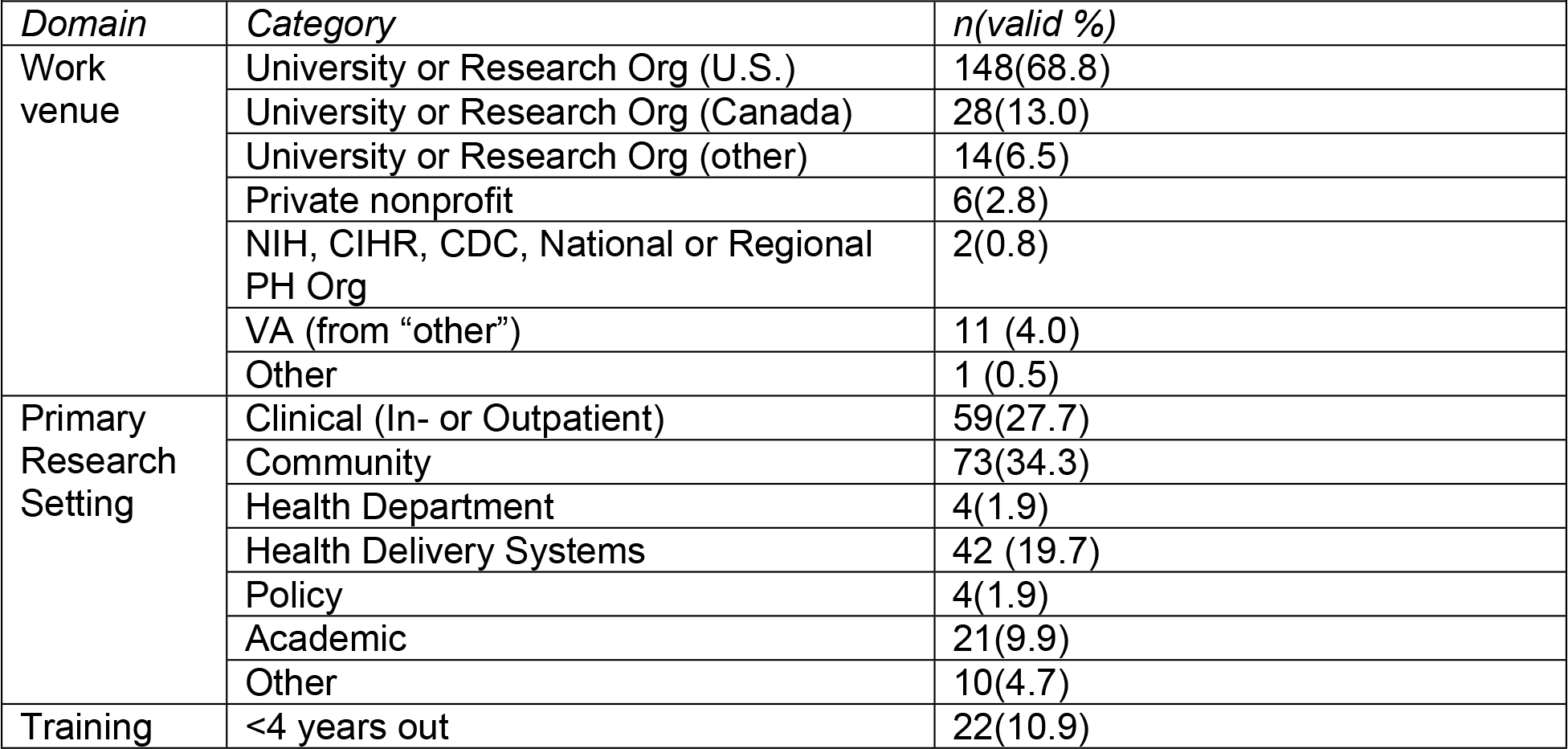
Characteristics of Survey Respondents

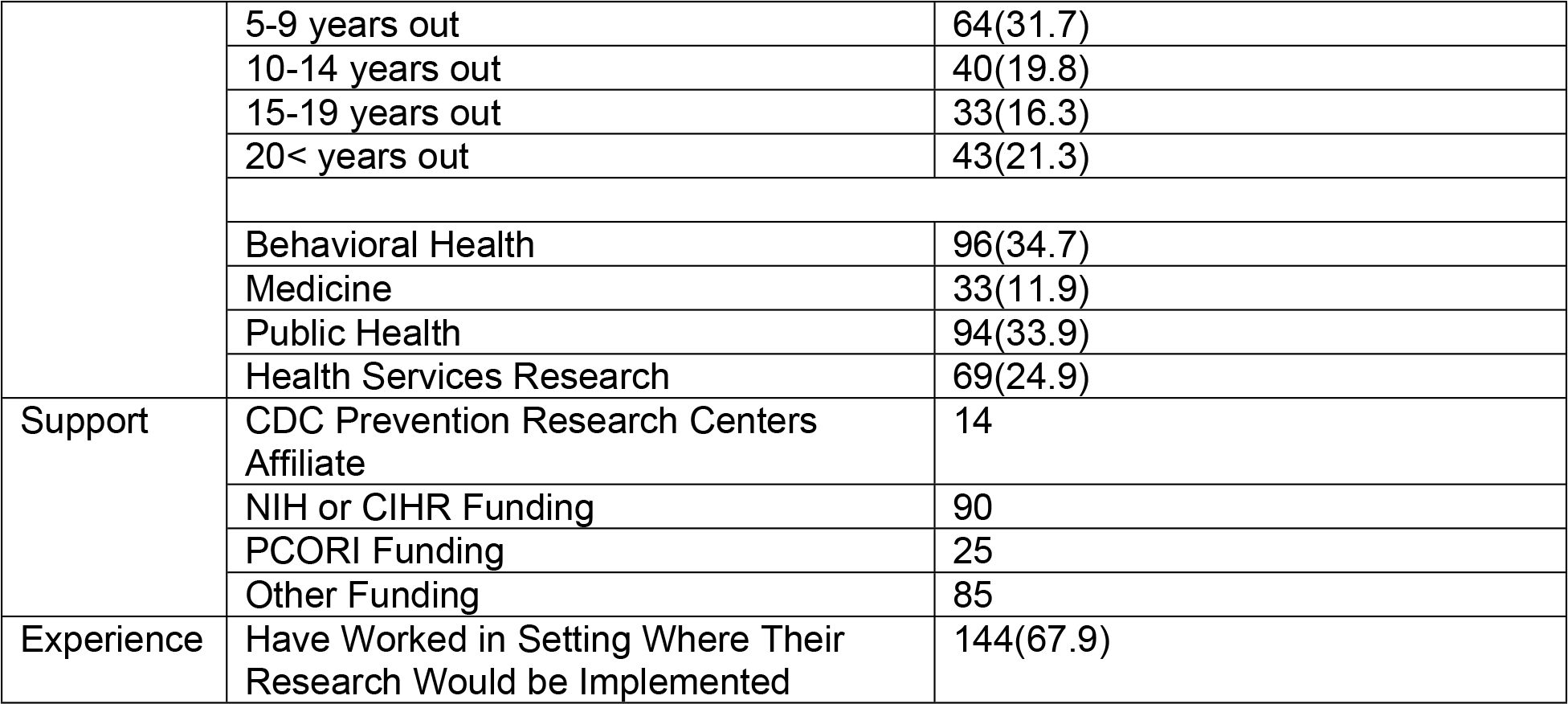

As seen in Table 2, the majority of respondents were from university or research settings in the U.S. (69%) or Canada (13%). They were from a mix of clinical (28%) and community settings (34%). The majority were from behavioral health (35%) or public health disciplines (34%); 26% had received formal training in D&I and there was a wide distribution in years since highest academic degree.

### Perceived Impact on Career and Practice/Policy

When prompted to respond with their level of agreement with the statement “It is an obligation of researchers to disseminate their research to those who need to learn about it and make use of the findings”, 56% of current respondents indicated that they strongly agree, compared to 51% in 2012. This difference was not statistically significant. When asked “how often do you involve stakeholders”, Brownson’s survey highlighted stakeholder engagement frequency at the project level, with 34% of participants saying they always or usually involved stakeholders, 49% sometimes or rarely, and 17% never. Individuals in our sample reported upon the frequency with which they typically engage non-research stakeholders *within* projects, with 55% of respondents indicating that they did so four or more times, 34% two to three times, 4% once, and 7% reported zero contacts with stakeholders.

### Reported Dissemination Activities

Respondents indicated routinely engaging in a variety of dissemination-related activities, with academic journals and conferences (88% in our survey vs. 86% in 2012 respectively), and reports to funders (74%, not included in 2012 survey) being the most frequent. Among these activities, publication in academic journals was identified as the most impactful on respondents’’ careers (94%), while face-to-face meetings with stakeholders are seen as most impactful on practice or policy (40%: Table 3).

**Table 3:** Dissemination Method & Priorities (all valid %)

| Table 3: Dissemination Method & Priorities (all valid %) |  |  |  |
| --- | --- | --- | --- |
|  | *Typically used* | *Most impact on career trajectory* | *Most impact on practice/policy* |
| Academic journals | 88 | 94 | 16 |
| Reports to funders | 74 | 0 | 6 |
| Press releases | 33 | 0 | 4 |
| Newsletters | 36 | 0 | 1 |
| Policy briefs | 21 | 0 | 8 |
| Social media | 42 | 0 | 3 |
| Academic conferences | 86 | 3 | 5 |
| Seminars/workshops | 51 | 1 | 9 |
| Face to face meetings with stakeholders | 55 | 0 | 40 |
| Media interviews | 22 | 0 | 1 |
| Webinars/videos | 30 | 0 | 0 |

### Stakeholder engagement

We asked several new questions related to stakeholder engagement in the 2018 survey. Stakeholder involvement in the research process was frequently reported, with clinical and community-based researchers engaging patients in similar ways (see Table 3) including focus groups and advisory committees. In terms of practitioner engagement, however, there were marked differences noted in that clinical researchers were more likely to include practitioners on user panels (47.5% vs. 24.4%, p<.01), as formal team members (62.7% vs. 45.5%, p <.05), and in interpreting data (59.3% vs. 38.2%, p<.05), but less likely than community researchers to use them in focus groups (43.9% vs. 10.5%, p<.01).

With respect to training, respondents who had more D&I training (defined as either a university course or fellowship) reported utilizing stakeholders differently than those with less (including workshops, other shorter trainings, or no formal D&I training). Those with more training were more likely to report using stakeholders in focus groups (including direct practitioners, organizational decision-makers, and policymakers). Similarly, those with more training were more likely to engage policymakers at all, being more likely to report engaging policymakers in focus groups, on advisory committees, as formal team members, and to help interpret data (Table 3).

Compared to their counterparts in the United States, university-based researchers in Canada reported engaging patients/consumers and direct practitioners in generally similar ways. However, Canadian researchers reported engaging organizational and policy-level stakeholders much more extensively than scientists in the U.S., with rate differences of greater than 25% observed between the two groups in terms of engagement of policymakers in focus groups (11.5 to 46%, p<.05), on advisory committees (33.1 to 61%, p<.05), on user panels (3.4 to 29%, p<.05), and as formal team members (8.8 to 43%, p<.05) (Table 4).

**Table 4:**
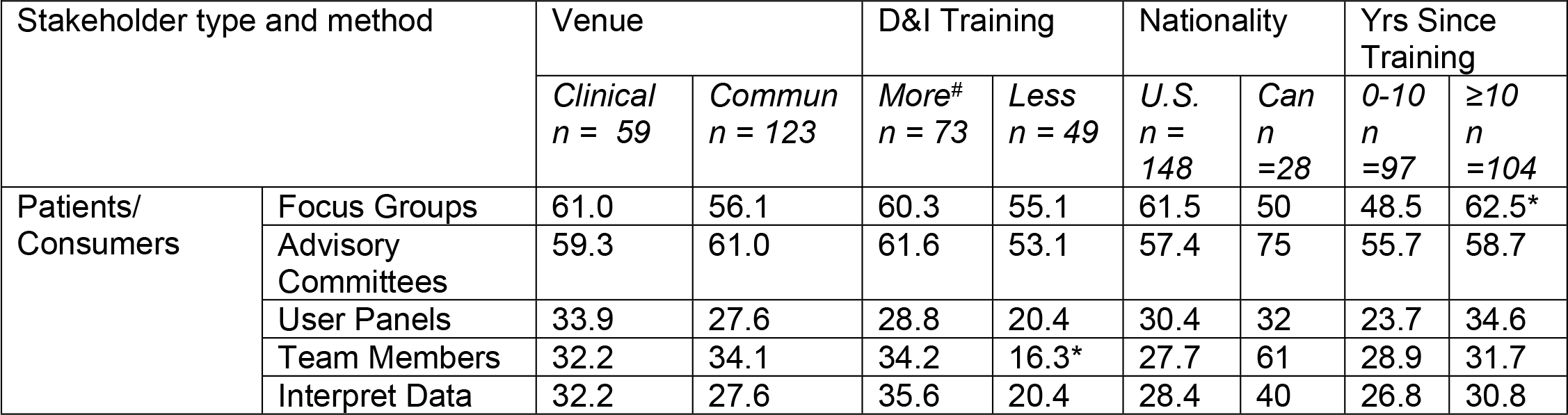
Stakeholder engagement type using various strategies (%)

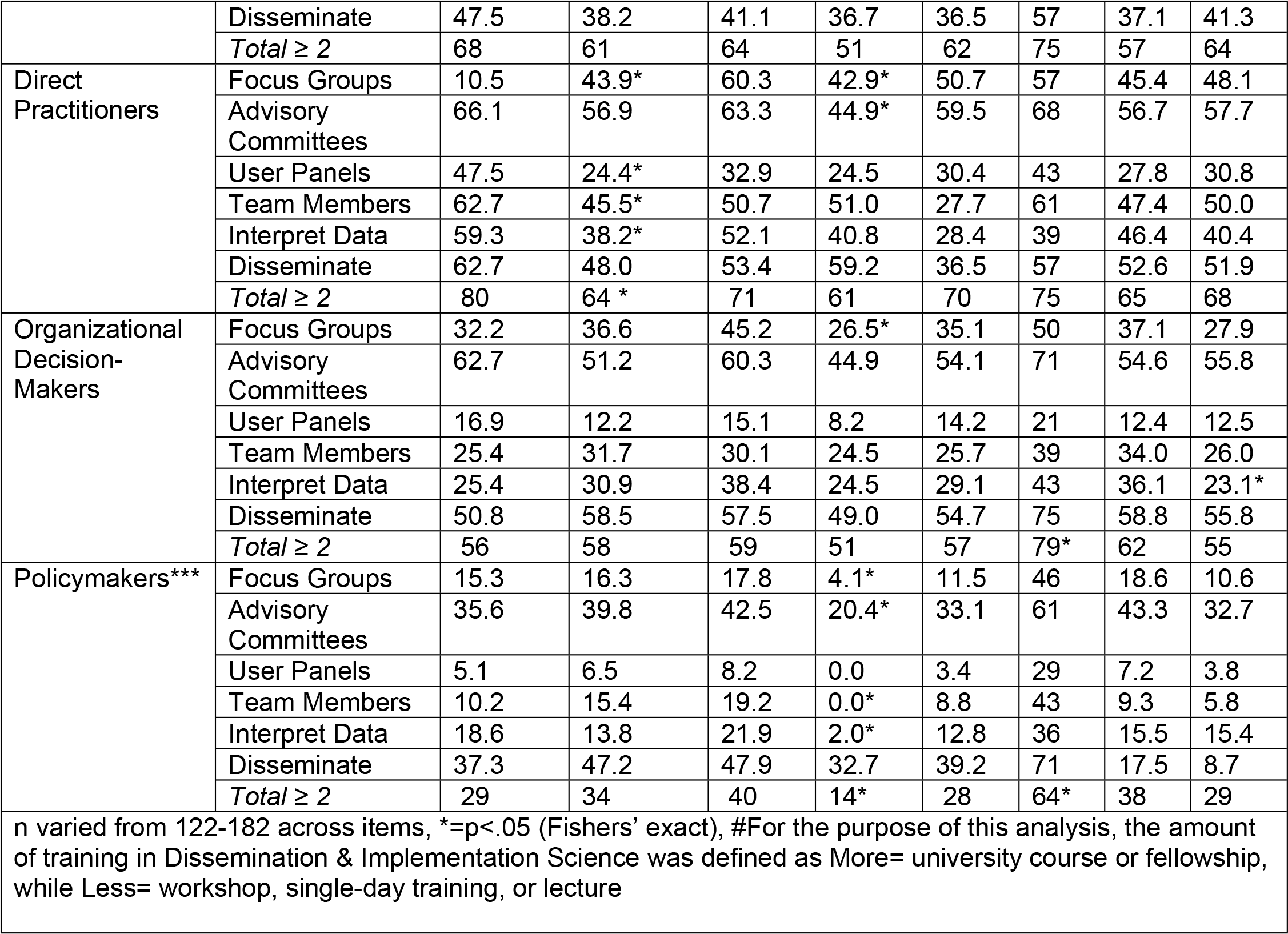

In response to a question regarding stage(s) within the research process during which stakeholders are involved, participants indicated that 28% involve them during the proposal phase (vs. 27% in Brownson et al.), 28% (vs.14% in 2011) in data gathering and analysis, and 47% (vs. 24%) in final reporting. Additional questions asked only in the present survey found that: 18% involved stakeholders across all research phases, 18% did not typically involve non-researchers, and 51% did so after publication for the purpose of supporting dissemination.

### Multivariable Analyses

Finally, logistic regression analyses were conducted to evaluate the relative contributions of different respondent characteristics to deeper or more comprehensive engagement of stakeholders. These analyses indicated that research venue, nationality, amount of D&I training, and seniority (assessed continuously as years since the respondent completed their graduate training) did not significant relate to routinely engaging patients, direct practitioners, or organizational decisionmakers in more than one way. None of these characteristics, significantly associated with an increased likelihood of engaging more than two stakeholder groups or routinely using more than the median number of total stakeholder engagement strategies (8).

## Discussion

With the increasing focus on disseminating research to practice (20) and a rapidly changing landscape of dissemination strategies, it is helpful to periodically assess what strategies D&I scientists are using to communicate evidence to practitioners and policy makers. This report updated and expanded the survey conducted in 2012 by Brownson and colleagues, but also sampled a broader range of D&I researchers (clinicians and Canadian D&I (KT) scientists), in addition to the public health researchers included in the 2012 sample, and provided greater depth on the evolving area of stakeholder engagement. D&I scientists reported engaging in varied dissemination activities, some but not all of which have increased, since the 2012 survey of public health researchers.

Comparisons were made on results on items that were identical or very similar to those reported in Brownson et al.’s earlier survey of public health researchers. Specifically, respondents in both samples reported using a variety of strategies to disseminate their work, but most frequently used traditional methods of publications in scientific journals and presentations at scientific meetings. This method is likely to influence the work of fellow researchers, who consistently report learning about emerging science in these venues (21, 22), but often neglects the seminars, professional association meetings, and electronic newsletters that local and state-level practitioners are more apt to use in their efforts to stay up-to-date (23). The ongoing predominance of these modes of dissemination today, despite recognition that these are not the most effective way to reach and engage practitioners, is likely due to the reward system of academic institutions. The general sentiment that dissemination of findings to non-research partners is a core responsibility of those engaged in academic pursuits appears to be shared between the two samples, despite several differences in the characteristics of the two samples.

Researchers in the 2018 sample still reported similar misalignment of incentives and behaviors related to dissemination of findings, as documented in the 2012 survey. One indication of the importance of reward structures can be seen in the details of the types and level of stakeholder engagement reported in the two samples. While the general rate of engaging stakeholders in research did not differ between the present survey and that reported by Brownson, the specific methods and depth of research engagement of stakeholders differed significantly between the two samples.

Even though both survey samples engaged stakeholders in similar ways, the stages at which stakeholders informed the research process differed. This was most apparent in the current sample, where 55% of respondents report that they typically engage stakeholders at least four times over the course of a project, whereas only 34% in Brownson’s survey reported that they typically involved stakeholders at all. We hypothesize that this increase may be due in part to the intervening impact of PCORI and other patient-centered funders requiring and stakeholder engagement for funding, although specific qualitative work is needed to understand these differences in greater detail.

### Differences across Respondent Types

Researchers in clinical and community settings, as well as those with more D&I training versus less, engage practitioners differently. Those with more D&I training were more likely to use a variety of stakeholder groups in a number of different ways. This was most evident as the stakeholder group engaged moved up the organizational or contextual scale: those with more D&I training were more likely to engage organizational and policy-level decision makers. In the latter case, there appears to be an important distinction in that these researchers appeared more likely to engage higher-level stakeholders at all, but were less likely to engage them in multiple ways. Future research and training should emphasize longitudinal involvement of higher-level stakeholders, including in varied roles.

Canadian researchers reported greater engagement, especially with policy makers. Those trained longer ago did not illustrate an appreciably different pattern of stakeholder engagement than those who completed their training more recently. Future analyses with higher power to detect subgroup differences should focus on hypothesized differences between these groups.

### Limitations and Future Directions

Although informative, this study has several limitations. These include the inability to determine a return rate given the unknown overlap among recruitment sources and the unknown number of researchers receiving invitations from listserv managers. Although we made concerted efforts to obtain representation from additional groups beyond those sampled in the 2012 survey, the limited number of respondents (and consequent insufficient statistical power to detect differences) in some categories such as Canadian researchers limit conclusions. Despite efforts to include a reasonable sample size of VA researchers, we were unable to obtain a sufficient number of VA researchers to conduct subgroup analyses.

The 2012 and 2018 sampling frames were purposively different and only a minority of the items directly replicated those on the 2012 survey. Other items were slight modifications and included additional dissemination response options that did not exist or were not applicable in 2012. As in any survey, our data are limited to respondents self-reported behavior and there may have been social demand characteristics to report, for example, greater levels of stakeholder engagement than are actually implemented. Despite these limitations, this study provided an important update on dissemination practices to non-research audiences and addressed a number of new questions such as the impact of D&I training on dissemination practices and assessment of the level and “depth” of stakeholder engagement practices.

Future directions include replication with larger samples and qualitative and mixed methods approaches to help understand some of the findings in greater depth. Gathering of data from a boader audience of scientists might yield more divergent use of stakeholder engagement and dissemination practices, theoretically yielding significant multivariable predictors of what makes a “high-quality disseminator”. Another question is how these and related findings (24) could buttress arguments for greater alignment between effective dissemination activities and academic incentives. This mis-alignment has persisted since the initial conference on dissemination and implementation (3), perpetuating a general heterogeneity of any dissemination efforts other than through traditional academic media. Finally, experimental comparisons of the actual effectiveness of different dissemination strategies on different outcomes (e.g., implementation of guidelines vs. policy change vs. de-implementation) are also indicated.

## Conclusions

Despite limited incentives for dissemination to non-research audiences, D&I researchers engage in a variety of strategies. There has been increased use of some, but not all strategies since 2012, and greater in depth and multi-level stakeholder engagement. Greater understanding of which dissemination strategies are most effective for what purposes and how to increase and sustain effective strategies is important to facilitate more rapid and complete translation of research to practice.

## List of Abbreviations

D&I: Dissemination and Implementation
NIH: National Institutes of Health
PCORI: Patient-Centered Outcomes Research Institute
CIHR KT: Canadian Institutes of Health Research Knowledge Translation
IRI: Implementation Research Institute.
MT-DIRC: Mentored Training for Dissemination and Implementation Research in Cancer
CDC: Centers for Disease Control and Prevention
VA QUERI: Veterans Association Quality Enhancement Research Initiative
KT Canada: Knowledge Translation Canada
NCI D&I: National Cancer Institute Dissemination and Implementation
TIDRH: Training Institute for Dissemination and Implementation Research in Health
Regional PH Org: Regional public health organization

## Declarations

### Ethics Approval & Consent to Participate

All activities described herein received prior approval from the Colorado Multiple Institutional Review Board (COMIRB).

### Consent for Publication

The authors consent to the full and unabridged publication of all manuscript and supplementary files.

### Availability of Data and Materials

The survey used in this study is included in this published article. The datasets used and/or analyzed during the current study are available on Zotero.

### Funding

Drs. Knoepke, Matlock, and Glasgow are supported the National Heart, Lung, and Blood Institute (R01HL136403). Dr. Knoepke is also supported by the American Heart Association (18CDA34110026). Dr. Brownson is supported by funding from the National Institute of Diabetes and Digestive and Kidney Diseases (P30DK092949, P30DK092950). The findings and conclusions in this article are those of the authors and do not necessarily represent the official positions of the National Institutes of Health or the American Heart Association. The funders had no role in study design, data collection and analysis, decision to publish, or preparation of the manuscript.

### Competing Interests

The authors declare that they have no competing interests

### Author Contributions

All authors contributed to the development of the survey. Survey responses were managed and catalogued by MPI. CK conducted all described statistical analyses. All authors reviewed, edited, and approved the final manuscript.

## Acknowledgements

We would like to acknowledge the support from the University of Colorado School of Medicine as well as the Dissemination & Implementation Program at the Adult & Child Consortium for Outcomes Research & Delivery Science (ACCORDS).

